# Stroke impairs the proactive control of dynamic balance during predictable treadmill accelerations

**DOI:** 10.1101/2024.10.18.618939

**Authors:** Tara Cornwell, James M. Finley

## Abstract

We maintain balance during gait using both proactive and reactive control strategies. Damage to the brain from a stroke impairs reactive balance, but little is known about how a stroke impacts proactive control during walking. Stroke-related impairments to proactive control could become targets for interventions designed to improve responses to predictable disturbances and reduce fall risk. Therefore, we determined if proactive strategies during predictable treadmill accelerations differed between people post-stroke (n=14) and people without stroke (n=14). Both groups walked with accelerations at random (every one to five strides) and regular (every three strides) intervals. We quantified the effects of the perturbations as changes to center of mass (COM) speed and used mechanical leg work to quantify the proactive strategies to slow the COM. Participants without stroke reduced peak COM speed better than those with stroke when perturbations were regular (−0.016 versus +0.004 m/s; p=0.007). They also reduced positive leg work more during the perturbation step than the group post-stroke (−5.7% versus +2.5%; p=0.003). One implication of these findings is that people post-stroke may be more susceptible to falls during predictable gait disturbances, and future work should identify the underlying impairments that cause these deficits.

## Introduction

People use a combination of proactive and reactive control strategies to maintain their balance while walking. Proactive control strategies generate motor commands to achieve a desired state based on knowledge about the body and the environment (1). When there are unexpected perturbations to a planned movement, deviations in the resultant state from the desired state, or errors, result in two coupled effects. Errors can trigger reactive control strategies and update the proactive mapping between error and reactive adjustments (1) through error-based learning. This learning process can be studied during walking with repeated perturbations, during which people use prior experience to reduce the destabilizing effects of the perturbation on their center of mass (COM) dynamics. After repeated practice, people reduce vertical displacement of the COM position during slips (2) and reduce trunk sway during both trips and slips (3). Identifying strategies to reduce the destabilizing effects of predictable perturbations can help us understand the roles of proactive versus reactive control during walking, the flexibility of our balance control, and how various impairments, whether due to age or injury, may affect fall risk.

Proactive strategies to reduce the destabilizing effects of an expected perturbation may include modifying foot placement or COM kinematics. When walking, the COM often moves outside the base of support (BOS), so people must reposition the COM within the BOS to prevent a possible fall (4). To do so, people may increase the distance between the COM and the edge of the BOS by adjusting foot placement or COM dynamics. For example, after repeated slips, people decrease step length and anteriorly shift their COM to reduce the chance of a backward loss of balance (5). A common measure of dynamic stability combines evaluations of the BOS and COM state in a single value known as the margin of stability (MOS) (6–9). The MOS is calculated as the distance between the edge of the BOS and one’s velocity-adjusted or extrapolated COM position (10). Taking longer steps or slowing the COM can each increase the MOS and configure the body to oppose a forward loss of balance, especially if the perturbation is predictable.

People may also proactively modify limb mechanics to reduce the destabilizing effects of the perturbation on the body. For example, reducing the positive work or increasing the negative work performed by the legs in anticipation of a forward loss of balance can each modify the magnitude or direction of the COM velocity. Most studies of walking employing predictable perturbations have not evaluated changes to kinetic metrics (3,5,6,9,11–14), but studies of running have reported that people decrease ground reaction forces during visible drops (15) or decrease leg stiffness on visible vertical steps (16,17). Thus, force-related adjustments to the control strategy may help mitigate the effects of an expected perturbation during walking.

While it is clear that adults without stroke make proactive adjustments after repeated exposure to perturbations, less is known about how a stroke affects the ability to make these adjustments. About one-third of chronic stroke survivors report experiencing at least one fall in a one-year period (18), but most research efforts to evaluate post-stroke control have been conducted during standing (19–22), and the limited number of studies in walking have mostly focused on reactive control with random perturbations (23–26). Various post-stroke impairments may hinder proactive adjustments during repeated perturbations. For example, cognitive deficits, observed in 60% of chronic patients (27), could hinder the ability to learn with experience. Muscle weakness and abnormal coordination (28) could hinder the ability to proactively modulate one’s gait pattern. Initial efforts to characterize post-stroke proactive control during walking have found that people post-stroke tripped over visible obstacles more frequently than their peers without stroke (12,29), perhaps because their stepping strategies were less flexible to the obstacle conditions (12). While obstacles help probe a visuomotor element of proactive control, obstacles are avoidable and thus may not be ideal for understanding the impact of movement errors or the reactive element of balance control. Instead, physical perturbations could more effectively examine the relative roles of proactive and reactive control strategies in people post-stroke.

Our goal was to investigate how the predictability of treadmill accelerations informs proactive and reactive balance control strategies in people with and without a stroke. Because people may learn the regular pattern, we hypothesized that regular accelerations would promote proactive changes to reduce the destabilizing effects on their COM dynamics. However, based on previous perturbation studies that found stepping strategies post-stroke less flexible (12) and less strongly correlated to COM state (30), we expected people post-stroke to maintain the same balance control strategies regardless of whether perturbations were predictable. This would make them less effective in reducing the destabilizing effects of the regular versus random perturbations. To test our hypotheses, we evaluated if people modulate fore-aft MOS and trailing and leading leg mechanical work to reduce the peak COM speeds during regular versus random accelerations and if these strategies differed in people post-stroke. Specifically, we expected that compared to random perturbations, regular perturbations would allow people without stroke to proactively increase their fore-aft MOS, perform less positive leg work, and perform more negative leg work to reduce peak COM speeds.

## Methods

### Participants

Fourteen adults with chronic stroke and 14 age-/sex-matched adults without stroke (Table 1) completed the following protocol. Participants were recruited through the University of Southern California Registry for Aging and Rehabilitation Evaluation (RARE) and the Healthy Minds Research Volunteer Registry. People post-stroke were recruited if they had a single stroke over six months prior and could walk unassisted for at least five minutes. People in either group were excluded if they had musculoskeletal conditions that limited walking ability, uncontrolled hypertension, or cognitive impairment (Montreal Cognitive Assessment score <19/30 (31)). Eight of the fourteen participants with a stroke reported using an ankle brace or orthotic (n=6), cane (n=2), or wheelchair (n=1) at least some of the time.

**Table 1.** Group means ± standard deviations of demographic characteristics and clinical scores. The last column displays p-values from the Mann-Whitney U-tests evaluating group differences. MoCA: Montreal Cognitive Assessment, Mini-BEST: Mini Balance Evaluation Systems Test, ABC Scale: Activities-specific Balance Confidence Scale.

|  | Non-stroke | Stroke | p-value |
| --- | --- | --- | --- |
| *Age (years)* | 58 ± 13 | 58 ± 13 | 0.945 |
| *Gender (M / F)* | 7 / 7 | 7 / 7 |  |
| *Height (m)* | 1.70 ± 0.10 | 1.66 ± 0.09 | 0.222 |
| *Mass (kg)* | 76 ± 14 | 75 ± 17 | 0.800 |
| *Treadmill speed (m/s)* | 1.05 ± 0.11 | 0.66 ± 0.17 | **<0.001** |
| *MoCA (0-30)* | 27 ± 2 | 26 ± 2 | 0.523 |
| *Mini-BEST (0-28)* | 26 ± 2 | 21 ± 4 | **<0.001** |
| *ABC Scale (%)* | 96 ± 5 | 78 ± 15 | **<0.001** |
| *Fugl-Meyer (0-34)* | | 23 ± 5 | |
| *Months since stroke* | | 109 ± 55 | |

### Experimental protocol

First, we performed several clinical assessments of motor impairment, cognition, and function. We used the Fugl-Meyer Assessment to measure the lower-extremity (LE) impairment post-stroke. For both groups, we administered the Montreal Cognitive Assessment (MoCA) to screen for cognitive impairment, Activities-specific Balance Confidence (ABC) scale to estimate self-reported confidence in one’s abilities to perform tasks without losing balance, Mini Balance Evaluations Systems Test (Mini-BEST) to evaluate dynamic balance, and 10-Meter Walk Test (10MWT) to calculate self-selected walking speeds (SSWS) overground. To determine participants’ SSWS on the treadmill, we used a staircase algorithm that started at 50% of their overground SSWS and stepped up or down according to their reported comfort.

Following the clinical assessments, participants completed eight walking trials on an instrumented dual-belt treadmill (Bertec, OH) at their SSWS. No handrails were provided to participants during any walking trial. During the two baseline trials, people acclimated to the treadmill (one minute) and then experienced three perturbations per side. A custom Python script triggered trip-like perturbations characterized by a trapezoidal speed profile that accelerated the belt at 5 m/s^2^ 250 ms before the predicted heel-strike to account for the delay in communication with the treadmill and held the belt speed at 150% SSWS for the duration of the estimated stance phase before decelerating back to SSWS at 5 m/s^2^ during the swing phase. The timing of the predicted heel strike and stance times were based on the median swing and stance times evaluated during the first 18 steps of the given trial, which consisted of unperturbed walking at SSWS. Based on pilot testing, the perturbation magnitude was scaled with walking speed to normalize the relative effects of the perturbation across participants. After the baseline trials, participants completed six perturbation trials, each consisting of 40 single-belt accelerations on the nondominant or paretic side at random (every one to five strides) or regular (every three strides) intervals. Perturbing one side helped maintain a consistent pattern for the regular interval block, and we chose the more impaired paretic side to better expose the effects of stroke. The order of the two blocks was randomized, and each was repeated three times before completing the other block of three trials. Performing three shorter trials per condition gave participants more frequent rest breaks to prevent fatigue. For safety, participants wore an overhead harness that did not provide body-weight support. They were also informed that perturbations would occur but were not provided information about the timing.

### Data acquisition and processing

Participants wore 54 reflective markers for 3D motion capture. We used Qualisys Track Manager to record the positions of these markers with a 10-camera optical motion capture system (Qualisys Oqus, Sweden) at 100 Hz and force data from the instrumented treadmill at 1000 Hz.

Data were then analyzed with custom Visual3D (HAS-Motion, Ontario, Canada) and MATLAB (MathWorks, Natick, MA) scripts. Marker data were low-pass filtered (Butterworth with 6-Hz cutoff) and gap-filled (third-order polynomial with a maximum gap of 10 frames). The body’s center of mass (COM) was estimated using a 13-segment (head, torso, upper arms, forearms, pelvis, thighs, shanks, and feet) model in Visual3D. We also identified and visually confirmed the following events in Visual3D: 1) heel strikes (HS) and toe offs as the maximum and minimum fore-aft positions of the heel and toe markers, respectively, 2) perturbation onset based on measures of the left and right belt speeds, and 3) “crossovers” when participants’ feet were placed on opposite belts. We removed an average of 17 steps per participant from force-related calculations because of crossover events. Additionally, we removed an average of 32 random-interval perturbation steps per participant because they occurred within one stride of the prior perturbation, as prior work found people with and without stroke require approximately two and three recovery steps, respectively, to return to a steady state after random treadmill accelerations (23).

### Peak COM speed as a measure of the destabilizing effects of the perturbations

Due to the trip-like nature of the treadmill accelerations, we expected to see increases in peak COM speed during perturbations (32) and reductions in this peak with predictable perturbations as people may try to reduce trajectory deviations for stability (33,34). We calculated COM speed as the magnitude of the COM velocity vector relative to the treadmill belt moving at SSWS during the perturbation steps. We considered all components of the velocity vector when computing speed because the perturbations elicit changes in both the anterior-posterior and vertical directions. Because walking speed affects peak speeds, we then normalized this metric to each participant’s SSWS. If perturbation predictability helps people effectively prepare, we would expect peak COM speeds to be slower during regular versus random accelerations.

### Fore-aft MOS

We evaluated the hypothesis that people would have a larger fore-aft margin of stability (MOS) upon HS of the regular perturbation step to reduce its destabilizing effects. Because the treadmill acceleration was applied in the sagittal plane, we expected that increasing the fore-aft MOS would be more helpful for counteracting the effects of the perturbation than modifying mediolateral MOS. We chose to evaluate MOS at HS because this would characterize the body state at the start of the perturbation and remove the influence of reactive adjustments made during the step. MOS was calculated as the distance between the edge of the base of support (BOS) and the extrapolated COM (10) in the fore-aft direction (Equation 1).

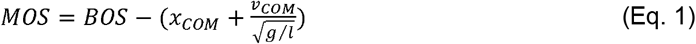

Here, we defined the leading edge of the BOS as the fore-aft position of the toe marker. The extrapolated COM is the COM position (*x*_*COM*_) adjusted by COM velocity (*v*_*COM*_), gravity (*g*), and leg length (*l*). Negative MOS values indicate that one’s extrapolated COM is outside the BOS, and a fall will occur unless one makes a corrective action, such as taking a step. People may prepare for predictable accelerations by increasing their fore-aft MOS at the HS of the perturbation step during the regular versus random conditions.

Mechanical leg work

The mechanical work performed by the leading and trailing legs can characterize another potential strategy employed in response to treadmill perturbations, as people modulate leg work depending on the perturbation’s timing (35). Mechanical power performed by the leg (*P*_*leg*_) was computed per step using the individual limbs method (36) as the sum of the power from the leg on the body and the power from the leg on the treadmill (Equation 2).

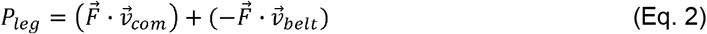

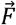 is the ground reaction force, 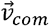 is the COM velocity, and 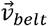 is the treadmill velocity relative to the lab coordinate system. We then computed work as the integral of power over time, with positive and negative work calculated separately to differentiate between energy generated and dissipated by the legs. Work was also normalized by subject mass to account for differences in weight between participants.

We quantified three potential changes to leg work that may reduce the destabilizing effects of predictable perturbations. First, people may decrease positive work performed by the trailing leg (Figure 1A) to reduce the initial forward COM speed at the beginning of the perturbation step and maintain the COM position further behind the leading edge of the BOS. The trailing leg takes the last contralateral step prior to the perturbation, so it will always refer to the non-paretic or dominant leg for the groups with and without stroke, respectively. Next, people may increase negative work performed by the leading leg during early stance (Figure 1B), absorbing some of the energy from the perturbation and reducing its destabilizing effects. Finally, people may decrease positive work performed by the leading leg (Figure 1B) for similar reasons as they may do so on the trailing leg. Thus, during regular versus random accelerations, we expected people to perform less positive work by the trailing and leading legs and more negative work by the leading leg.

**Figure 1.**
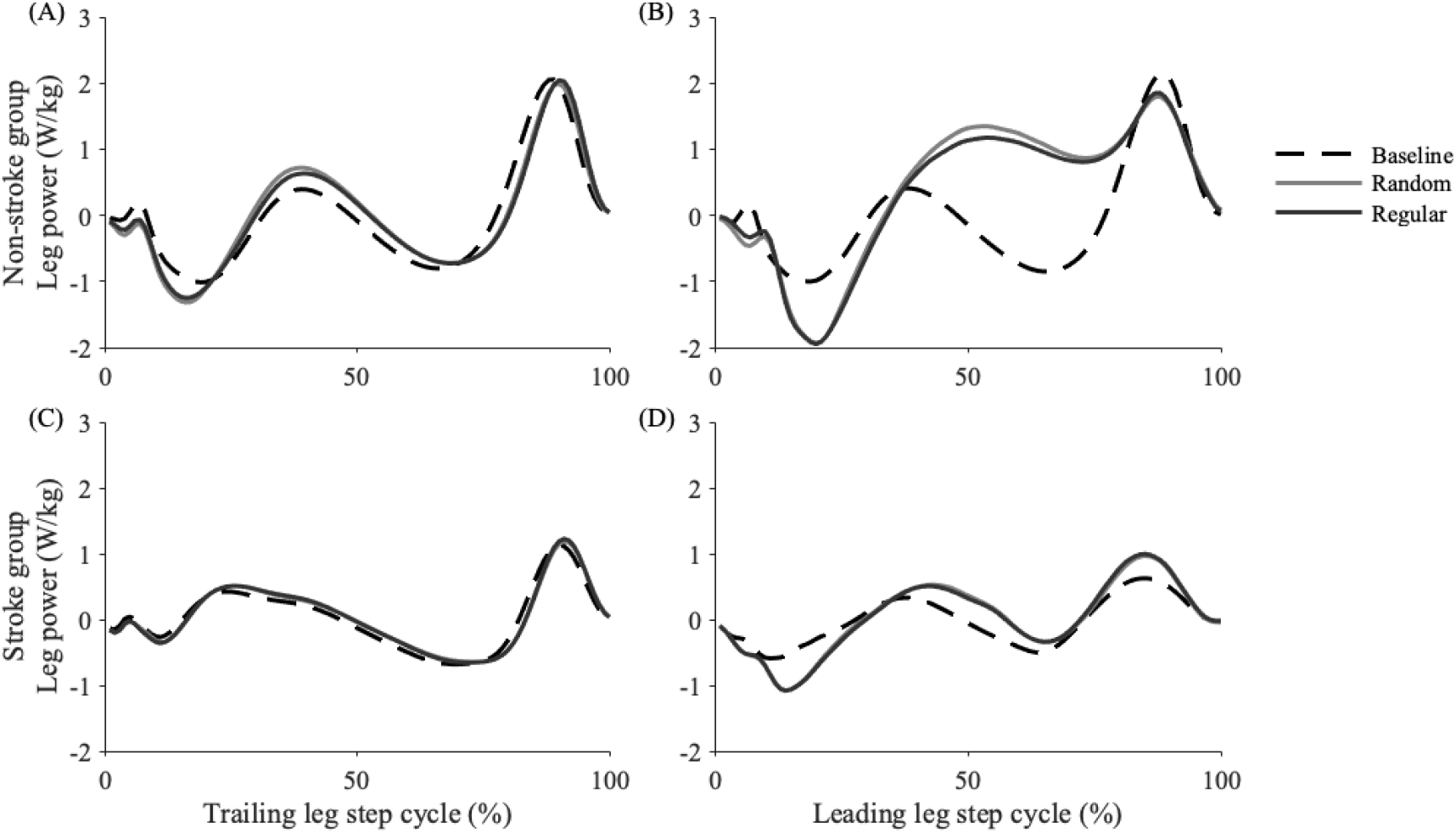
Mean mechanical leg power performed by the trailing and leading legs for the (A-B) non-stroke and (C-D) stroke groups. Power is plotted across the step cycle during the baseline, unperturbed condition (dashed), random-interval condition (gray solid), and regular-interval condition (black solid). The baseline condition includes all steps, but only pre-perturbation (trailing leg) and perturbation (leading leg) steps were considered for the random- and regular-interval conditions. While there were minimal visible changes to trailing leg power in either group (A, C), the perturbations elicited changes to leading leg power with more energy absorption (negative) early in stance but more energy generation (positive) starting around midstance in the group without stroke (B).

### Statistics

Statistical analyses were performed in MATLAB version R2022b, and alpha was set to 0.05. Our primary aim was to understand how each of our outcome measures varied between groups and trials. We used separate linear mixed-effects models to evaluate the effects of Group (two categorical levels), Trial (two categorical levels), and the interaction between Group and Trial on each normalized variable. We hypothesized that people post-stroke would have smaller reductions in peak COM speed than those without stroke during regular versus random perturbations. We also evaluated potential proactive strategies by quantifying changes in fore-aft MOS and mechanical leg work. Random intercepts accounted for between-subject differences in baseline values. We plotted standardized residuals versus fitted values to check for heteroscedasticity and nonlinearity. One outlier with a standardized residual greater than three was removed from the leading leg positive work model, and the model was then refit.

Finally, we were interested in determining if measures of motor or cognitive impairment were associated with one’s ability to reduce the destabilizing effects of predictable perturbations. Therefore, we used linear regressions to evaluate associations between clinical measures (SSWS, Mini-BEST, Fugl-Meyer, and MoCA) and the change in peak COM speed, our primary outcome measure. We calculated leverage values and removed two high scores for SSWS, representing outliers, before refitting the model.

## Results

### Participants without stroke mitigated the destabilizing effects of regular accelerations

As intended, our treadmill perturbations caused changes in whole-body kinematics. The destabilizing nature of the accelerations is demonstrated by a representative participant without stroke whose average peak COM speeds increased during perturbed versus unperturbed steps (Figure 2). For additional context, during random accelerations, participants with and without stroke reached average peak COM speeds of 0.86 and 1.28 m/s, 0.05 and 0.09 m/s faster, respectively, than those experienced during unperturbed baseline walking.

**Figure 2.**
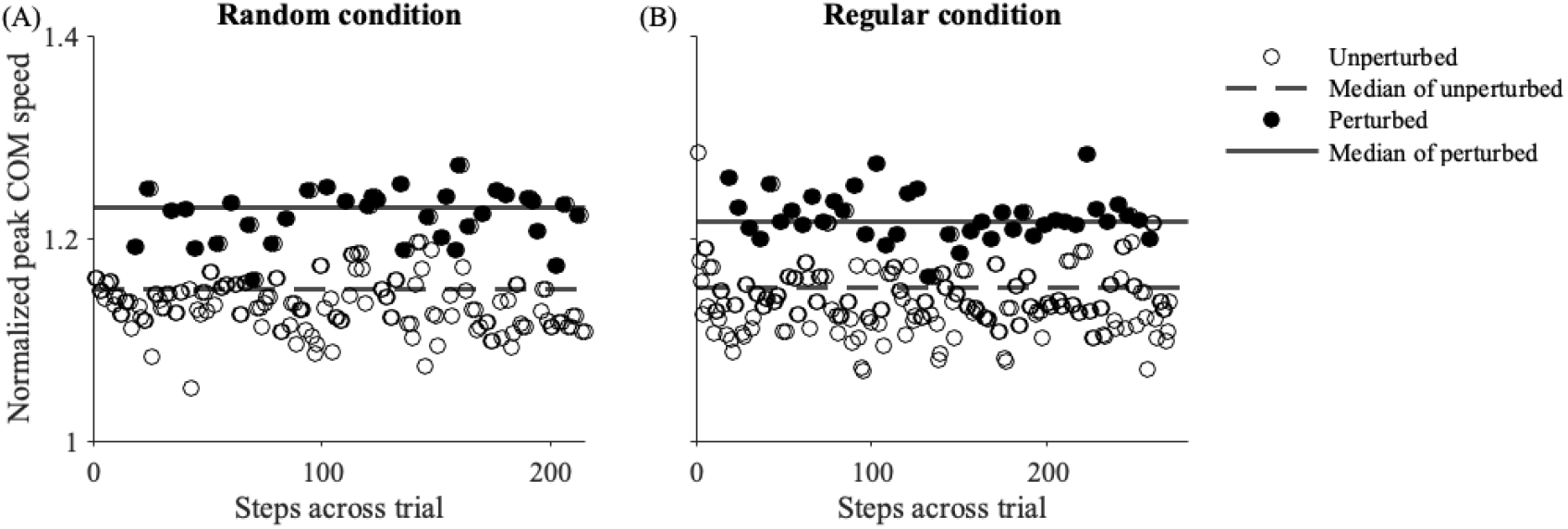
Representative data from a participant without stroke. Peak center of mass (COM) speeds normalized by self-selected walking speed are plotted per step during the final trials with (A) random- and (B) regular-interval accelerations. Normalized peak COM speeds during unperturbed steps and their median values are plotted in open circles and dashed lines, while peak COM speeds during perturbed steps and their median values are plotted in black circles and solid lines.

When normalized by SSWS, peak COM speeds during perturbations were slower in the group without stroke than those with stroke (Figure 3A; Group p=0.001). Compared to the random-interval condition, participants used the predictability from the regular intervals to slow peak COM speeds (Figure 3A; Trial p=0.017), and this reduction was greatest in the group without stroke (Figure 3B; Group*Trial p=0.007). Participants without stroke best mitigated the destabilizing effects of the regular versus random perturbations, with a change in peak COM speed of -0.016 m/s (−1.2%) versus +0.004 m/s (+0.6%) in the group post-stroke.

**Figure 3.**
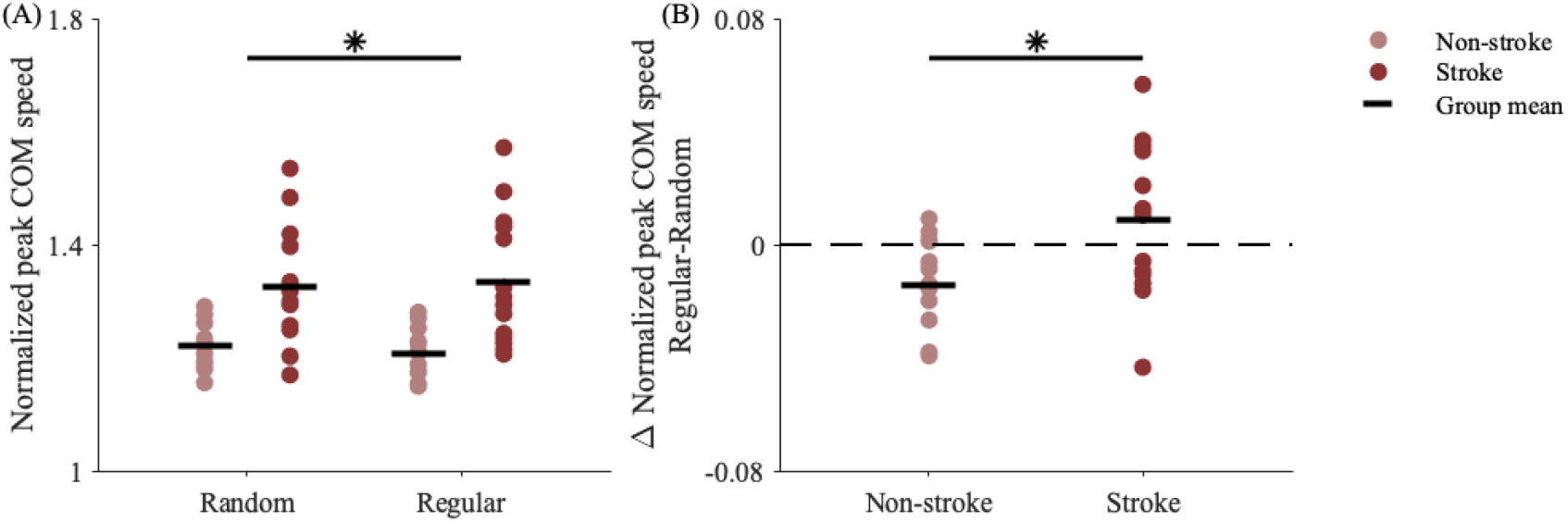
(A) Peak center of mass (COM) speeds normalized by self-selected walking speeds. Normalized peak center of mass (COM) speeds were slower in the group without versus with stroke (Group p=0.001) and during regular versus random accelerations (Trial p=0.017). (B) Changes in normalized peak COM speeds during regular versus random accelerations. The group without stroke had a significantly greater reduction in peak COM speed than the group with stroke during regular versus random accelerations (Group*Trial p=0.007).

### People proactively increase fore-aft margin of stability with regular perturbations

We next evaluated several potential proactive changes that people may make to reduce the destabilizing effects of the perturbation, including changes to the body’s configuration at initial contact and changes in mechanical work before or during the perturbation. People increased the fore-aft MOS at HS of regular versus random accelerations (Figure 4) with no significant group differences within trials (Trial p=0.010; Group*Trial p=0.266). However, the magnitudes of these changes were relatively small, with average increases of 0.4 ± 0.5 cm and 0.9 ± 0.3 cm in the groups with and without stroke, compared to unperturbed MOS values of 16.2 ± 4.1 cm and 14.5 ± 2.7 cm, respectively.

**Figure 4.**
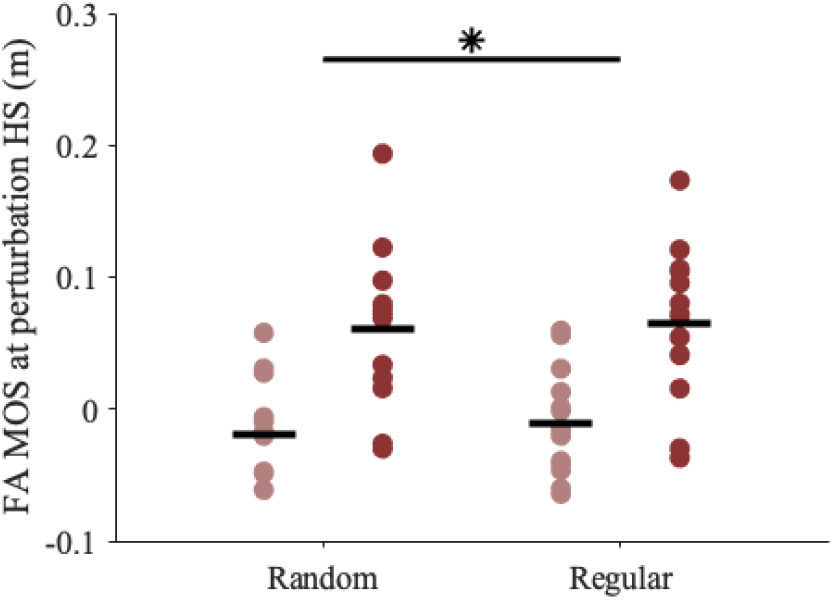
Fore-aft (FA) margin of stability (MOS) at heel-strike (HS) of the perturbation step was significantly larger during the regular-versus random-interval trials (Trial p=0.010). There was no significant group difference within trials (Group*Trial p=0.266).

### People without stroke perform less positive leg work with regular perturbations

Another proactive strategy could involve modulating mechanical leg work during the pre-perturbation or perturbation steps to mitigate the effects of the predictable perturbation. However, people did not modify positive work performed by the trailing leg (Trial p=0.075; Group*Trial p=0.289) or negative work performed by the leading leg (Trial p=0.079; Group*Trial p=0.216). Instead, people, particularly those without stroke, proactively performed less positive work by the leading leg (Figure 5A) during regular versus random accelerations (Trial p<0.001; Group*Trial p=0.003). Participants without stroke performed an average of 0.02 J/kg (5.7%) less positive work by the leading leg during regular versus random accelerations (Figure 5B). This positive work typically occurred during midstance (Figure 1B), and reducing this work would also help slow the COM. The significant interactions between Group and Trial suggest people post-stroke did not demonstrate the same adjustments to leading leg positive work during regular accelerations as their peers without stroke, which may have hindered their ability to reduce the destabilizing effects of the predictable perturbations.

**Figure 5.**
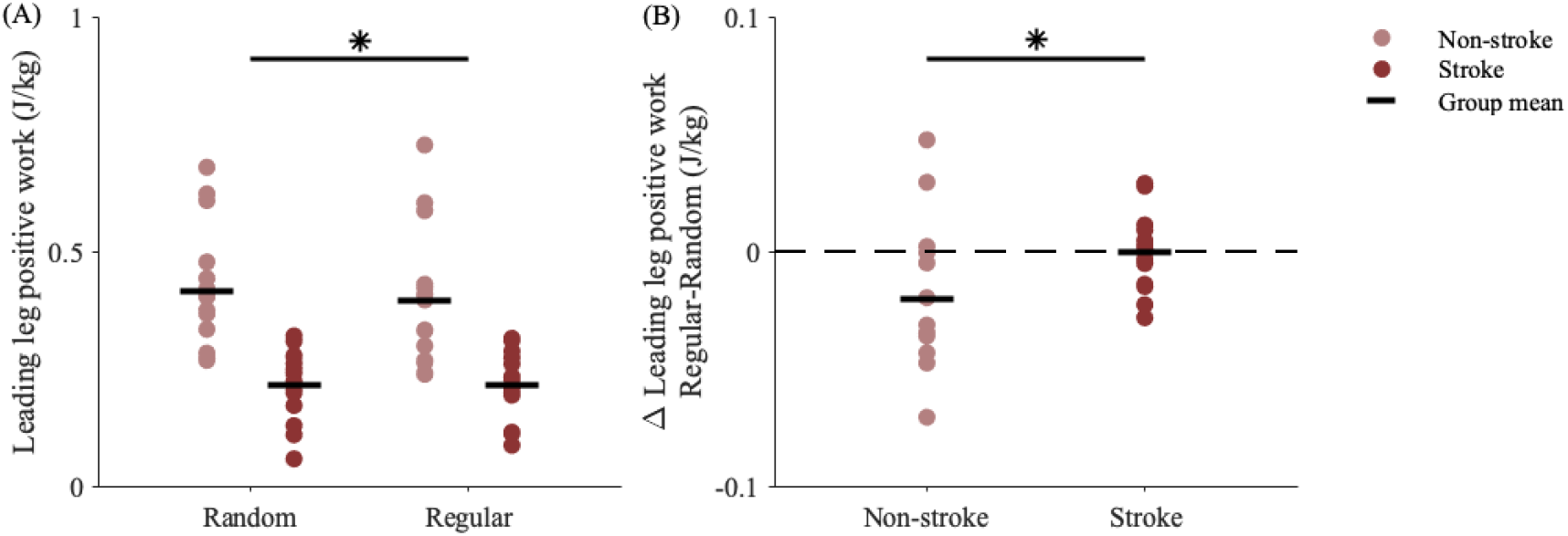
(A) Positive work performed by the leading leg in each group during random and regular perturbation steps. (B) Change in leading leg positive work (Regular-Random). Compared to the group post-stroke, the group without stroke performed less positive work with the leading leg during regular versus random perturbations (Group*Trial p=0.003).

To elucidate which proactive changes were associated with the observed reductions in peak COM speeds during regular accelerations, we fit a multiple linear regression model with the percent changes in MOS at perturbation step HS and leading leg positive work as the two predictors of the percent change in peak COM speeds during regular versus random accelerations. After confirming non-collinearity among the predictors and removing four outliers identified by high leverage values, we found that the change in leading leg positive work, but not fore-aft MOS (p=0.400), was a significant predictor of the change in peak COM speeds. Then, we refit the model (excluding one outlier) with only the statistically significant predictor and found that performing less positive work by the leading leg explained approximately 20% of the reductions in peak COM speeds during regular versus random accelerations (R^2^=0.223, p=0.004).

### Change in perturbation response is correlated with SSWS

We used exploratory linear regression models to evaluate the relationships between clinical measures (SSWS, Mini-BEST, Fugl-Meyer, and MoCA) and the observed differences in peak COM speeds during regular versus random accelerations. Interestingly, changes in COM speed were not correlated to Mini-BEST, Fugl-Meyer, or MoCA scores but were negatively correlated with SSWS (Figure 6). Participants who walked faster, which may indicate higher motor function, experienced slower peak COM speeds (R^2^=0.317, p=0.002) during regular versus random accelerations.

**Figure 6.**
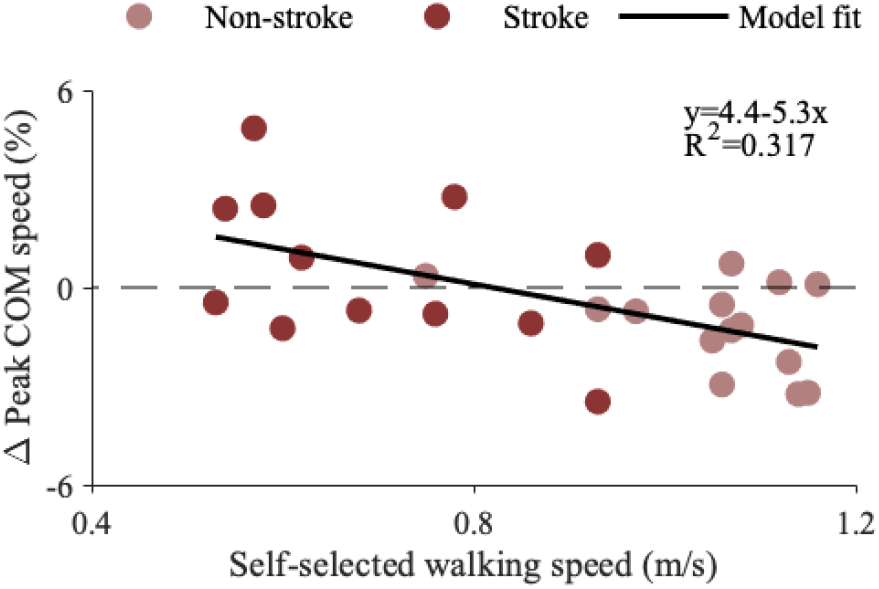
Linear regression model relating self-selected walking speed with the percent changes in peak center of mass (COM) speed during regular versus random perturbations (R^2^=0.317, p=0.002). Individual means are plotted (circles) with the model fit line (solid black). Group was not considered in the model. Data from the two people post-stroke who walked at the slowest speeds were identified as outliers based on high leverage values and removed. If these two outliers are included, walking speed is still a significant predictor of the changes in peak COM speeds (R^2^=0.328, p<0.001).

## Discussion

Here, we investigated the effects of regular versus random treadmill accelerations on balance control strategies during walking and how these strategies are affected by stroke. We quantified changes in peak COM speeds to evaluate the effects of perturbation predictability. Then, we evaluated possible proactive strategies with measures of fore-aft MOS and mechanical leg work. In support of our first hypothesis, participants without stroke were less disturbed by perturbations with regular versus random intervals, suggesting that predictability informs effective proactive control strategies.

### Balance control strategies differ in response to random versus predictable disturbances

Our participants without stroke performed less positive leg work during accelerations at regular versus random intervals, and these changes explained over 20% of the reductions in peak COM speed. Our findings indicating that people make proactive adjustments to leg work, combined with prior work demonstrating that people modify ground reaction forces and leg stiffness when running with expected terrain changes (15–17), suggest that modifying the force produced by the leg may be an important and energy-efficient (37) control strategy for people without neurologic injury.

People may also make other proactive adjustments, but there were no significant changes in the trailing leg positive work or leading leg negative work between conditions. While there was a significant increase in fore-aft MOS on the heel strike of regular versus random perturbations, the difference was small and did not significantly predict the changes observed in peak COM speeds. Therefore, our findings suggest that people do not adjust their body state or steps in preparation for an expected perturbation. This aligns with previous work that did not observe proactive adjustments in young adults, who, in response to auditory warnings about impending treadmill belt accelerations, did not change MOS or whole-body angular momentum (13), and guinea fowl, who did not adjust their step timing when treadmill obstacles were presented at slower speeds that allow for more anticipatory control (38). Overall, adjustments to leg work, not foot placement, seemed to help our participants without stroke mitigate the destabilizing effects of regular accelerations, demonstrating how implicit knowledge of a perturbation’s timing can inform and update control plans during walking.

### Post-stroke impairments to proactive balance control and their clinical implications

Proactive control adjustments may become more important in populations with greater fall risk or impaired reactive responses, such as older adults (39,40) and people post-stroke (18,23). However, taking advantage of perturbation predictability may also become more challenging, as seen in people post-stroke during standing perturbations (19–22) and obstacle negotiation tasks (12,29). To our knowledge, this is the first study to evaluate how the regularity of physical perturbations informs balance control during walking and how this process is affected by a stroke. In support of our second hypothesis, we found that people post-stroke were less flexible in modulating control strategies and less effective in improving stability with perturbation regularity than their peers without stroke.

During regular versus random treadmill accelerations, people post-stroke slowed their peak COM speeds less than their peers. There was also a correlation between these changes and walking speed (Figure 6), underlining the impact of physical impairment on balance control. Additionally, people post-stroke did not demonstrate the same modifications to work performed by the leading leg, nor any compensatory changes to work performed by the trailing leg. The leading or perturbed leg was always the paretic, normally weaker side, which may help explain the control deficits seen on this side. For example, sensory deficits (41) could reduce or delay the sensory feedback from the foot experiencing the acceleration, or abnormal muscle coordination (28) could prevent the muscle-or joint-level changes necessary to modulate leg work. Previous work also found proactive control deficits emphasized on the paretic side during predictable standing perturbations, during which participants stopped a swinging pendulum with their outreached palms (22). The participants without stroke demonstrated anticipatory muscle activity in most of the eight leg and trunk muscles measured, but the group post-stroke only had anticipatory paretic-side activity in the tibialis anterior before the perturbation (22). Our results complement these findings, suggesting a stroke also impairs proactive control during walking, though this should also be tested on the non-paretic side with bilateral perturbations. While our goal was to determine if impairments in proactive control were associated with clinical measures, we did not find associations between the observed changes in peak COM speeds and Mini-BEST, lower extremity Fugl-Meyer, or MoCA scores, highlighting the difficulty in identifying the root cause of individual deficits. Future studies should further investigate proactive control post-stroke with different perturbations, predictability modalities, and clinical tests of motor and cognitive deficits to determine the underlying impairments that limit proactive adjustments.

Improving walking ability and balance is a major rehabilitation goal post-stroke, so the search for effective clinical tools and protocols is ongoing. Recently, perturbation training has emerged as a potential tool to practice balance strategies, with some promising initial results in standing and walking. Following a two-week training protocol with predictable standing perturbations, people post-stroke demonstrated earlier anticipatory muscle activations and smaller COM displacements from the perturbation (42), indicating a more effective control strategy. Training studies during walking have mostly implemented random perturbations (24–26). However, practice with predictable perturbations in a safe environment may help identify effective and achievable proactive strategies, build strength and coordination, and instill balance confidence. Our results suggest that people post-stroke exhibit control deficits during predictable gait perturbations, so future work should evaluate whether these control strategies are adaptable and transferrable to real-world conditions. Training protocols with repeated, predictable perturbations should test the effects of training on balance control responses under novel conditions and future falls in daily living. By practicing both proactive and reactive elements of balance control, people post-stroke may be able to safely improve their strategies and reduce their fall risk.

### Limitations

One of the limitations of the current study arises from the group differences in walking speed, which may confound the effects of a stroke. While matching the speeds between the groups could have alleviated this concern, it would have required participants without stroke to walk much slower than they would naturally, and their lack of familiarity with walking at this speed would introduce a new confound to the results. Instead, we chose to normalize the size of the perturbation to walking speed and normalize peak COM speed by walking speed to evaluate relative changes between conditions. Because the magnitude of the speed changes was smaller for those walking at slower speeds, it is possible that the participants walking at slower speeds, most of whom were in the group post-stroke, did not perceive the perturbations as threatening enough to their balance to necessitate proactive changes to control. While we did not directly assess the perceived threat of the perturbations, our participants post-stroke experienced greater relative increases in COM speed than participants without stroke. In addition, prior research has shown people with stroke were significantly more destabilized than people without stroke when responding to treadmill accelerations that increased speeds by 0.2 m/s, which is smaller than the perturbations our participants experienced (23). Therefore, although there was a significant difference in speeds between the two groups, our findings, combined with those from prior work, suggest that the applied perturbations were moderately disturbing for our participants post-stroke, such that group differences in proactive control may be due to participant-specific sensorimotor impairments.

Additionally, while we evaluated several potential proactive strategies, there may be other key proactive strategies that we did not fully characterize. For example, by understanding joint- and muscle-level contributions to proactive control strategies, we could identify post-stroke impairments such as delayed or reduced muscle activation and potentially target these impairments through rehabilitation interventions such as biofeedback-based training. Using perturbation predictability to “set” neuron excitability and scale reflexive muscle responses could lead to faster recovery from expected balance disturbances (43,44). Therefore, central set could be assessed by evaluating changes in muscle activity in response to perturbations (2,45). Lastly, modulating the mechanical impedance of the leg could help mitigate the effects of a predictable perturbation. Impedance quantifies the resistance of a joint or limb to displacement in response to an applied load and can be modulated by changes in muscle activation before a perturbation, through reflexes, or voluntarily after a perturbation (46–49). Future studies should use a combination of electromyography and joint- or limb-level perturbations to evaluate the proactive regulation of reflex sensitivity and changes in limb impedance during gait perturbations.

## Conclusions

We varied the regularity of treadmill accelerations to elicit changes to proactive balance control strategies. Compared to people post-stroke, people without stroke better mitigated the destabilizing effects of regular versus random perturbations, perhaps by modifying leading leg mechanical work. Walking speed was correlated with changes in peak COM speeds between conditions, suggesting those with greater motor impairment are less able to modify proactive strategies during predictable gait perturbations and may be more susceptible to falls. Future work should examine a broader set of potential proactive strategies and determine if these strategies are modifiable with practice, particularly in individuals post-stroke.

## Acknowledgment

This work was supported by the American Heart Association Predoctoral Fellowship 23PRE1012432, the Eunice Kennedy Shriver National Institute of Child Health & Human Development of the National Institutes of Health under award number R01HD091184, and NSF award number 2319710. We would also like to thank all participants for their significant contributions to our work.

## Data availability

Data and code used in this project can be found at https://osf.io/t8ue4/.

## Notes

https://osf.io/t8ue4/

### Competing Interest Statement

The authors have declared no competing interest.

### Summary of Updates

Analyses were simplified; Discussion section was updated to expand on limitations

## References

1. Kandel ER, editor. Principles of neural science. 5th ed. New York: McGraw-Hill; 2013. 743–767 p.

2. Marigold DS, Patla AE. Strategies for Dynamic Stability During Locomotion on a Slippery Surface: Effects of Prior Experience and Knowledge. J Neurophysiol. 2002 Jul;88(1):339–53.

3. Brodie MA, Okubo Y, Sturnieks DL, Lord SR. Optimizing successful balance recovery from unexpected trips and slips. J Biomech Sci Eng. 2018;13(4):17–00558.

4. Patla AE. Strategies for dynamic stability during adaptive human locomotion. IEEE Eng Med Biol Mag. 2003 Mar;22(2):48–52.

5. Bhatt T, Wening JD, Pai YC. Adaptive control of gait stability in reducing slip-related backward loss of balance. Exp Brain Res. 2006 Mar 1;170(1):61–73.

6. Hak L, Houdijk H, Steenbrink F, Mert A, van der Wurff P, Beek PJ, et al. Speeding up or slowing down?: Gait adaptations to preserve gait stability in response to balance perturbations. Gait Posture. 2012 Jun 1;36(2):260–4.

7. McAndrew Young PM, Wilken JM, Dingwell JB. Dynamic margins of stability during human walking in destabilizing environments. J Biomech. 2012 Apr 5;45(6):1053–9.

8. Roeles S, Rowe PJ, Bruijn SM, Childs CR, Tarfali GD, Steenbrink F, et al. Gait stability in response to platform, belt, and sensory perturbations in young and older adults. Med Biol Eng Comput. 2018 Dec 1;56(12):2325–35.

9. Major MJ, Serba CK, Chen X, Reimold N, Ndubuisi-Obi F, Gordon KE. Proactive Locomotor Adjustments Are Specific to Perturbation Uncertainty in Below-Knee Prosthesis Users. Sci Rep. 2018 Jan 30;8(1):1863.

10. Hof AL, Gazendam MGJ, Sinke WE. The condition for dynamic stability. J Biomech. 2005 Jan;38(1):1–8.

11. Wang TY, Bhatt T, Yang F, Pai YC. Adaptive control reduces trip-induced forward gait instability among young adults. J Biomech. 2012 Apr 30;45(7):1169–75.

12. Den Otter AR, Geurts ACH, de Haart M, Mulder T, Duysens J. Step characteristics during obstacle avoidance in hemiplegic stroke. Exp Brain Res. 2005 Feb 1;161(2):180–92.

13. Eichenlaub EK, Urrego DD, Sapovadia S, Allen J, Mercer VS, Crenshaw JR, et al. Susceptibility to walking balance perturbations in young adults is largely unaffected by anticipation. Hum Mov Sci. 2023 Jun 1;89:103070.

14. Kreter N, Lybbert C, Gordon KE, Fino PC. The effects of physical and temporal certainty on human locomotion with discrete underfoot perturbations. J Exp Biol. 2022 Oct 13;225(19):jeb244509.

15. Müller R, Ernst M, Blickhan R. Leg adjustments during running across visible and camouflaged incidental changes in ground level. J Exp Biol. 2012 Sep 1;215(17):3072–9.

16. Ferris DP, Liang K, Farley CT. Runners adjust leg stiffness for their first step on a new running surface. J Biomech. 1999 Aug 1;32(8):787–94.

17. Grimmer S, Ernst M, Günther M, Blickhan R. Running on uneven ground: leg adjustment to vertical steps and self-stability. J Exp Biol. 2008 Sep 15;211(18):2989–3000.

18. Schmid AA, Yaggi HK, Burrus N, McClain V, Austin C, Ferguson J, et al. Circumstances and consequences of falls among people with chronic stroke. 2013;50(9):1277–86.

19. Marigold DS, Eng JJ, Timothy Inglis J. Modulation of ankle muscle postural reflexes in stroke: influence of weight-bearing load. Clin Neurophysiol. 2004 Dec 1;115(12):2789–97.

20. Marigold DS, Eng JJ. Altered timing of postural reflexes contributes to falling in persons with chronic stroke. Exp Brain Res. 2006 Jun 1;171(4):459–68.

21. Joshi M, Patel P, Bhatt T. Reactive balance to unanticipated trip-like perturbations: a treadmill-based study examining effect of aging and stroke on fall risk. Int Biomech. 2018 Jan 1;5(1):75–87.

22. Curuk E, Lee Y, Aruin AS. Individuals With Stroke Use Asymmetrical Anticipatory Postural Adjustments When Counteracting External Perturbations. 2019 Oct 1;23(4):461–71.

23. Liu C, McNitt-Gray JL, Finley JM. Impairments in the mechanical effectiveness of reactive balance control strategies during walking in people post-stroke. Front Neurol. 2022;13.

24. Matjačić Z, Zadravec M, Olenšek A. Feasibility of robot-based perturbed-balance training during treadmill walking in a high-functioning chronic stroke subject: a case-control study. J NeuroEngineering Rehabil. 2018 Apr 11;15(1):32.

25. Handelzalts S, Kenner-Furman M, Gray G, Soroker N, Shani G, Melzer I. Effects of Perturbation-Based Balance Training in Subacute Persons With Stroke: A Randomized Controlled Trial. Neurorehabil Neural Repair. 2019 Mar 1;33(3):213–24.

26. Esmaeili V, Juneau A, Dyer JO, Lamontagne A, Kairy D, Bouyer L, et al. Intense and unpredictable perturbations during gait training improve dynamic balance abilities in chronic hemiparetic individuals: a randomized controlled pilot trial. J NeuroEngineering Rehabil. 2020 Jun 17;17(1):79.

27. Nakling AE, Aarsland D, Næss H, Wollschlaeger D, Fladby T, Hofstad H, et al. Cognitive Deficits in Chronic Stroke Patients: Neuropsychological Assessment, Depression, and Self-Reports. Dement Geriatr Cogn Disord Extra. 2017 Aug 29;7(2):283–96.

28. Arene N, Hidler J. Understanding motor impairment in the paretic lower limb after a stroke: a review of the literature. Top Stroke Rehabil. 2009;16(5):346–56.

29. van Swigchem R, van Duijnhoven Hjr, den Boer J, Geurts AC, Weerdesteyn V. Deficits in Motor Response to Avoid Sudden Obstacles During Gait in Functional Walkers Poststroke. Neurorehabil Neural Repair. 2013 Mar 1;27(3):230–9.

30. Dragunas AC, Cornwell T, López-Rosado R, Gordon KE. Post-Stroke Adaptation of Lateral Foot Placement Coordination in Variable Environments. IEEE Trans Neural Syst Rehabil Eng. 2021;29:731–9.

31. Trzepacz PT, Hochstetler H, Wang S, Walker B, Saykin AJ, for the Alzheimer’s Disease Neuroimaging Initiative. Relationship between the Montreal Cognitive Assessment and Mini-mental State Examination for assessment of mild cognitive impairment in older adults. BMC Geriatr. 2015 Sep 7;15(1):107.

32. Lee-Confer JS, Finley JM, Kulig K, Powers CM. Reactive responses of the arms increase the Margins of Stability and decrease center of mass dynamics during a slip perturbation. J Biomech. 2023 Aug 1;157:111737.

33. Hicheur H, Pham QC, Arechavaleta G, Laumond JP, Berthoz A. The formation of trajectories during goal-oriented locomotion in humans. I. A stereotyped behaviour. Eur J Neurosci. 2007;26(8):2376–90.

34. Bucklin MA, Wu M, Brown G, Gordon KE. American Society of Biomechanics Journal of Biomechanics Award 2018: Adaptive motor planning of center-of-mass trajectory during goal-directed walking in novel environments. J Biomech. 2019 Sep 20;94:5–12.

35. Golyski PR, Sawicki GS. Which lower limb joints compensate for destabilizing energy during walking in humans>? J R Soc Interface. 2022 Jun;19(191):20220024.

36. Selgrade BP, Thajchayapong M, Lee GE, Toney ME, Chang YH. Changes in mechanical work during neural adaptation to asymmetric locomotion. J Exp Biol. 2017 Aug 15;220(16):2993–3000.

37. Donelan JM, Kram R, Kuo AD. Mechanical work for step-to-step transitions is a major determinant of the metabolic cost of human walking. J Exp Biol. 2002 Dec 1;205(23):3717–27.

38. Gordon JC, Rankin JW, Daley MA. How do treadmill speed and terrain visibility influence neuromuscular control of guinea fowl locomotion>? J Exp Biol. 2015 Oct 1;218(19):3010–22.

39. Blake AJ, Morgan K, Bendall MJ, Dallosso H, Ebrahim SBJ, Arie THD, et al. Falls by elderly people at home: Prevalence and associated factors. Age Ageing. 1988 Jan 1;17(6):365–72.

40. Tang PF, Woollacott MH. Inefficient Postural Responses to Unexpected Slips During Walking in Older Adults. J Gerontol Ser A. 1998 Nov 1;53A(6):M471–80.

41. Sullivan JE, Hedman LD. Sensory Dysfunction Following Stroke: Incidence, Significance, Examination, and Intervention. Top Stroke Rehabil. 2008 May 1;15(3):200–17.

42. Curuk E, Aruin AS. Perturbation-based training enhances anticipatory postural control in individuals with chronic stroke: a pilot study. Int J Rehabil Res. 2022 Mar;45(1):72–8.

43. Evarts EV, Shinoda Y, Wise SP. Neurophysiological approaches to higher brain functions. 1. print. New York, N.Y.: Wiley; 1984. 198 p. (A Neurosciences Institute publication).

44. Prochazka A. Sensorimotor gain control: A basic strategy of motor systems>? Prog Neurobiol. 1989 Jan;33(4):281–307.

45. Horak FB, Diener HC, Nashner LM. Influence of central set on human postural responses. J Neurophysiol. 1989 Oct;62(4):841–53.

46. Krutky MA, Ravichandran VJ, Trumbower RD, Perreault EJ. Interactions Between Limb and Environmental Mechanics Influence Stretch Reflex Sensitivity in the Human Arm. J Neurophysiol. 2010 Jan;103(1):429–40.

47. Krutky MA, Trumbower RD, Perreault EJ. Influence of environmental stability on the regulation of end-point impedance during the maintenance of arm posture. J Neurophysiol. 2013 Feb 15;109(4):1045–54.

48. Rouse EJ, Hargrove LJ, Perreault EJ, Kuiken TA. Estimation of Human Ankle Impedance During the Stance Phase of Walking. IEEE Trans Neural Syst Rehabil Eng. 2014 Jul;22(4):870–8.

49. Jakubowski KL, Ludvig D, Bujnowski D, Lee SSM, Perreault EJ. Simultaneous Quantification of Ankle, Muscle, and Tendon Impedance in Humans. IEEE Trans Biomed Eng. 2022 Dec;69(12):3657–66.

